# Comprehensive analysis of the Aquaporin genes in *Eucalyptus grandis* suggests potential targets for drought stress tolerance

**DOI:** 10.1101/2023.10.29.564209

**Authors:** Dayana S. Seidel, Paulo H. Claudino, Gabriela Sperotto, Simone N. Wendt, Zachery D. Shomo, Ravi V. Mural, Henrique M. Dias

## Abstract

This study delves into the comprehensive analysis of *AQP* genes in *Eucalyptus grandis*, providing insights into their genomic abundance, diversification, expression patterns across tissues, and responses to drought stress. We identified 53 *AQP* genes in the *Eucalyptus grandis* genome, categorized into four subfamilies: *AQP-NIP*, *AQP-SIP*, *AQP-PIP*, and *AQP-TIP*. This abundance of *AQP* genes is a reflection of gene duplications, both tandem and whole-genome, which have shaped their expansion. The chromosomal distribution of these genes reveals their widespread presence across the genome, with some subfamilies exhibiting more tandem duplications, suggesting distinct roles and evolutionary pressures. Sequence analysis uncovered characteristic motifs specific to different AQP subfamilies, demonstrating the diversification of protein and targeting. The expression profiles of *AQP* genes in various tissues in both *Arabidopsis thaliana* and *Eucalyptus grandis* showcased variations, with root tissues showing higher expression levels. Notably, *AQP-PIP* genes consistently exhibited robust expression across tissues, highlighting their importance in maintaining water regulation within plants. Furthermore, the study investigated the response of *AQP* genes to drought stress and rehydration, revealing differential expression patterns. *EgAQP-NIP* and *EgAQP-TIP* genes were up-regulated during drought stress, emphasizing their role in osmotic equilibrium and water transport. Conversely, *EgAQP-PIP* genes showed down-regulation during drought stress but were up-regulated upon rehydration, indicating their involvement in water movement across cell membranes. Overall, this research contributes to our understanding of *AQP* genes in *Eucalyptus grandis*, shedding light on their genomic evolution, expression patterns, and responses to environmental challenges, particularly drought stress. This information can be valuable for future studies aimed at enhancing the drought resilience of woody perennial plants like *Eucalyptus grandis*.

## INTRODUCTION

The Aquaporins (AQPs) have been extensively investigated using genome sequencing across numerous plant species as they play a vital role in the regulation of water. Specifically, AQPs have been extensively researched in rice (*Oryza sativa*) (Sakurai *et al*. 2005), soybean (*Glycine max*) (Zhang *et al*. 2013), cotton (*Gossypium L.*) (Li *et al*. 2019), grape (*Vitis vinifera*) (Fouquet *et al*. 2008), and poplar (*Populus trichocarpa*) (Gupta and Sankararamakrishnan 2009). The aquaporin superfamily comprises membrane-bound proteins that contain six characteristic membrane-spanning alpha helices and can range from 26 to 34 kDa (Tyerman *et al*. 2002). Also known as membrane intrinsic proteins (MIPs), the first plant aquaporin to be purified was from *Glycine max* and was annotated to be NODULIN- 26 (GmNOD26) (Fortin *et al*. 1987). After decades of additional studies on structure, transport, and regulatory mechanisms, it became evident that in addition to water transport, AQPs are also essential for pore formation and metabolite trafficking across membranes from one aqueous space to another (Meli *et al*. 2018).

As the AQPs are found in a variety of cell types and organelles, understanding their structure and function as they relate to water and metabolite transport has proven to be a crucial step in improving crop quality and tolerance to abiotic and biotic stresses (Zenda *et al*. 2023). Prior research has shown that AQPs in vascular plants are able to aid in physiological and developmental processes, including cell and tissue expansion, fiber development, signal transduction during pollination, and seed development by facilitating movement of soluble compounds across membranes (Azad *et al*. 2004; Eisenbarth and Weig 2005; Gattolin *et al*. 2011; Soto *et al*. 2008, 2010; Vander Willigen *et al*. 2006; Wudick *et al*. 2014). In plants, many specialized AQPs aid in this necessary metabolic shunting throughout the cell. The plasma membrane intrinsic proteins (PIPs), allow for water transfer across the plasma membrane. Other AQPs such as the tonoplast intrinsic proteins (TIPs), nonulin26-like intrinsic proteins (NIPs), and small basic intrinsic proteins (SIPs) have more diverse roles including the transport of water, metabolites, and other small molecules (Maurel 2007).

In addition to their unique functions, the AQPs also display tissue-specific and developmental expression patterns (Javot and Maurel 2002). This allows for targeted regulation of water and solute transport during critical developmental stages and in response to environmental changes (Alexandersson *et al*. 2005). Given the diversity in both function and subcellular location of the AQPs, there is a growing interest in targeting them as candidates to improve plant resilience (Altpeter *et al*. 2005). Studies on AQPs and response to abiotic stress have already begun to correlate AQPs with increased tolerance to drought, salinity, cold, and heat (Kapilan *et al*. 2018; Wang *et al*. 2020). While there is much interest in the AQPs of crop species, work has also focused on the roles AQPs play in woody plants such as *Eucalyptus grandis* and their relationship with osmotic and water stress (Rodrigues *et al*. 2013, 2016; Feltrim *et al*. 2021).

By combining techniques in genomics, transcriptomics, proteomics, and metabolomics, we can gain insights into the genetic and molecular mechanisms that underlie the ability of woody plants to withstand environmental pressures. This presents a promising avenue for improving the stress tolerance of these species. Therefore, given the unique challenges associated with studying the genetics of forest plants, collaboration among experts in plant physiology, genetics, ecology, and environmental science can enable a comprehensive examination of woody plants from multiple perspectives (Lobo *et al*. 2022; Rowland *et al*. 2023).

Here, we investigate the biological aspects of AQPs in *Eucalyptus grandis* using a comparative genomics approach. We elucidate the diversity of paralogs compared to *Arabidopsis thaliana* along with variations in protein motifs and expression patterns of the AQPs across different tissues. While many of the AQP genes in *Eucalyptus grandis* have been characterized based on their sequence, the biological roles of many AQPs remain unknown. We have identified 53 AQP homologs in *Eucalyptus grandis* and analyzed their phylogeny, motif compositions, and expression patterns using a comprehensive bioinformatics approach. Taken together, our work provides several vignettes into the expression patterns, dynamics, and evolutionary trends of the AQPs and lays the foundation for further research into AQPs in *Eucalyptus grandis*.

## MATERIAL AND METHODS

### Database Search and Sequence Mining

The search and identification of *AQP* genes in *Eucalyptus grandis* was performed in the PLAZA Eudicot 5.0 database (Van Bel *et al*. 2018) (https://bioinformatics.psb.ugent.be/plaza/versions/plaza_v4_5_monocots/), using the AQP gene sequence of *Arabidopsis thaliana* (*AtAQP* - AT4G10380) as a query (Ward 2001). The orthologous gene family of *AQP* was defined in PLAZA Eudicot 5.0 under accession number ORTHO05D001001 (https://bioinformatics.psb.ugent.be/plazaversions/plaza_v4_5_monocots/). All putative candidates were verified using the online software InterProScan (Blum *et al*. 2021) and Pfam (El-Gebali *et al*. 2019) to check the presence of the full MIP protein family domain (IPR000425).

### Multiple Alignments and Phylogenetic Analysis

Protein alignments were performed using the MAFFT v7 web tool (Katoh *et al*. 2019) (https://mafft.cbrc.jp/alignment/ server/), following default analysis parameters (global alignment; BLOSUM62 score matrix; and Gaps penalty = 1.5). The tree topology was defined using the PhyML v3 tool (Guindon *et al*. 2010) (http://www.atgc-montpellier.fr/phyml/), wherein the best model was predicted by "SMS: Smart Model Selection” (Lefort *et al*. 2017), the branch support was determined by “aLRT SH-like” model, and the phylogenetic tree was structured using the “iTOL” web tool (https://itol.embl.de).

### Gene and Protein Features Analysis

The physical positions of the *AQP* gene sequence of *Eucalyptus grandis* (*EgAQP*) on the chromosome were visualized using the MG2C web tool () (Chao *et al*. 2021). In addition, collinearity analysis was performed with CoGe (Lyons and Freeling 2008; Lyons *et al*. 2008) (https://genomevolution.org/coge/SynMap.pl) for determination of duplication events and substitution non-synonymous (Kn), synonymous (Ks) and ratio Kn/Ks values in pairwise comparison between *Eucalyptus grandis* (Myburg *et al*. 2014) and *E. urophyla x grandis* (Shen *et al*. 2023). Furthermore, the conserved protein motifs were investigated with MEME suite (Bailey *et al*. 2009) (http://meme.nbcr.net). The subcellular localization prediction was performed using the Plant-mPloc web tool (Chou and Shen 2010). Alignments were visualized with WebLogo (http://weblogo.berkeley.edu/logo.cgi) (Crooks *et al*. 2004)), to generate graphical representations (logos) of the NPA-box patterns within a multiple sequence alignment.

### *In Silico* Expression Analysis

RNAseq data were obtained from independent experiments and research groups on the NCBI database (https://www.ncbi.nlm.nih.gov). The RNAseq datasets for different tissues of *Arabidopsis thaliana* were assigned with the accession numbers PRJNA194429, PRJNA242915, PRJNA195608, PRJNA151589, PRJNA231089 and PRJNA234023, while *Eucalyptus grandis* datasets were under accession number PRJNA253676. Specifically, the RNAseq dataset was chosen for the drought experiment in *Eucalyptus grandis* with accession PRJNA896601. All datasets underwent a uniform analysis pipeline, which included quality assessment of the reads using FastQC, adapter trimming with Trim Galore!, and mapping and quantification via Salmon quant. Transcript assemblies of *Arabidopsis thaliana* and *Eucalyptus grandis* were employed to conduct these analyses, available in the PLAZA database (Van Bel *et al*. 2018) (Dicots 4.5; https://bioinformatics.psb.ugent.be/plaza/versions/plaza_v4_5_monocots/). All analysis pipelines were performed with default parameters on the Galaxy Europe analysis platform (https://usegalaxy.eu). Mean expression values were extracted in transcripts per million (TPM) for each experimental condition along with their corresponding standard deviations.

## RESULTS

### Identification and Classification of AQP Protein Superfamily Members in *Eucalyptus grandis*

To explore putative *EgAQP* homologs in *Eucalyptus grandis*, the sequence AT4G10380 corresponding to an *AtNIP5;1* was used as the query on the Plaza database. The AQP family was automatically annotated and received an ortho family number accession (ORTHO05D001001). A total of 53 *AQP* homologs were identified in the *Eucalyptus grandis* genome (**Table 1**). The homologs were manually curated by considering their domain structure and phylogenetic protein analysis. The resulting clusters were categorized into four AQP subfamilies, as described in Diehn *et al*. (2015). 48 AQP proteins carrying entire MIP domain were subsequently classified into subfamilies AQP-NIP, AQP-SIP, AQP-PIP, and AQP-TIP, based on a comparative analysis (**Figure 1**; **Supplementary Table S1**) and a review of previous studies in Eucalyptus (Rodrigues *et al*. 2013, 2016; Feltrim *et al*. 2021).

**Figure 1.**
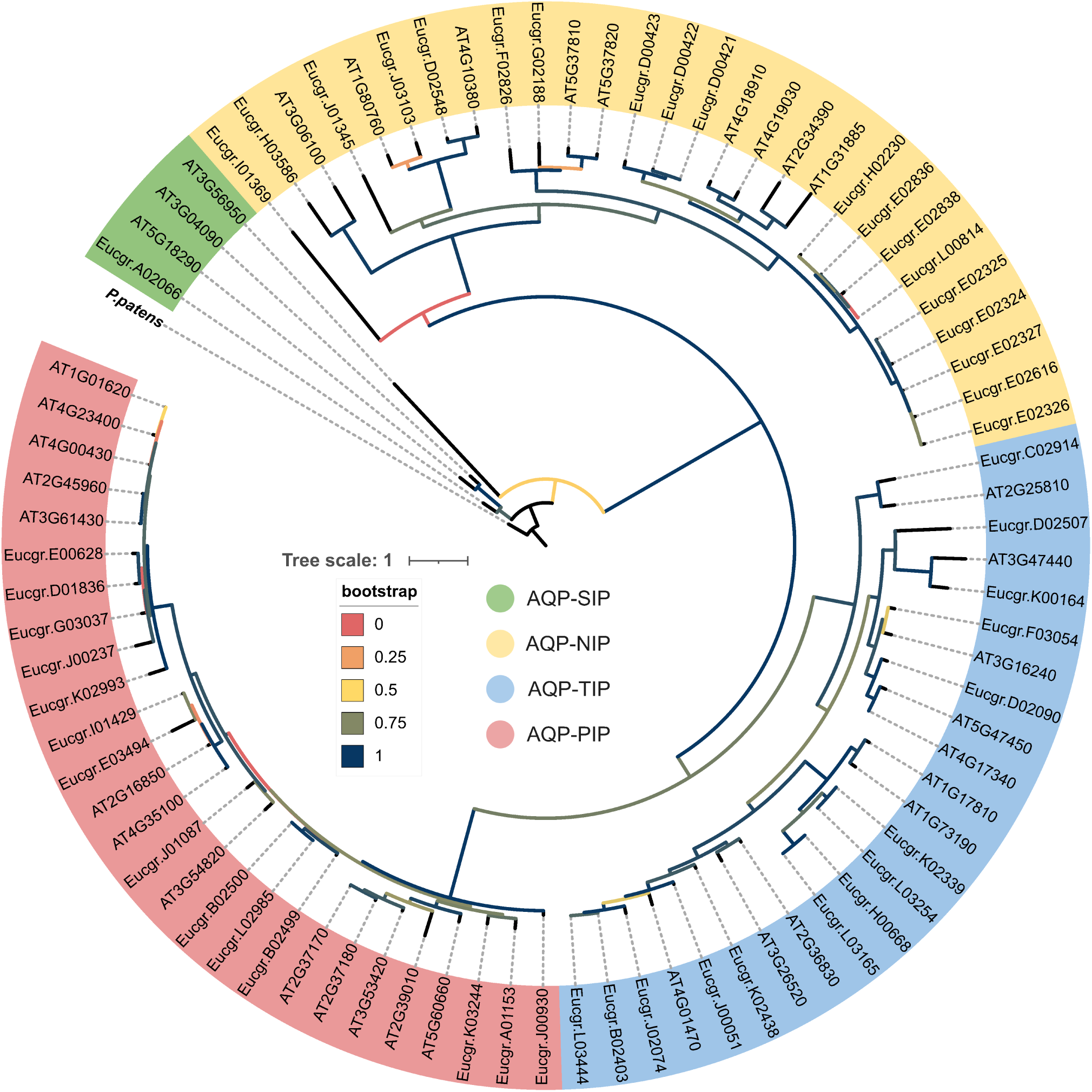
The sequences of AQPs from *Eucalyptus grandis* underwent a global multiple alignment constructed using MAFFT v7. This alignment highlights the annotation of the Major Intrinsic Protein (MIP) domain across all proteins and low conservation degrees along protein homologs. It also underscores the conservation of the NPA-box residues within the AQPs.

**Table 1.** Identification and classification of AQP genes in the *Eucalyptus grandis* and *Arabidopsis thaliana* genomes. Nomenclature, clade, strand, location, and subcellular localization prediction are presented.

| Clade | Symbol | Gene ID | Species | Strand | Chr | Location | Subcellular location prediction (mPloc) |
| --- | --- | --- | --- | --- | --- | --- | --- |
| I | AtAQP-SIP | AT1G14230 | At | + | Chr1 | 4861497-4863991 |  |
| I | AtAQP-SIP | AT1G14240 | At | + | Chr1 | 4865159-4867777 |  |
| I | AtAQP-SIP | AT1G14250 | At | + | Chr1 | 4868675-4871203 |  |
| I | AtAQP-SIP | AT2G02970 | At | - | Chr2 | 865395-868007 |  |
| I | AtAQP-SIP | AT3G04080 | At | - | Chr3 | 1068068-1070917 |  |
| I | AtAQP-SIP | AT3G04090 | At | - | Chr3 | 1072340-1074031 |  |
| I | AtAQP-SIP | AT3G56950 | At | - | Chr3 | 21078586-21079644 |  |
| I | AtAQP-SIP | AT5G18280 | At | - | Chr5 | 6050799-6054023 |  |
| I | AtAQP-SIP | AT5G18290 | At | - | Chr5 | 6055198-6056407 |  |
| I | EgAQP-SIP | Eucgr.A02066 | Eg | + | Chr01 | 36162345-36165428 | Cell membrane |
| I | EgAQP-SIP | Eucgr.A02068 | Eg | + | Chr01 | 36169338-36172690 | Nucleus |
| I | EgAQP-SIP | Eucgr.A02176 | Eg | + | Chr01 | 37070264-37073242 | Cell membrane |
| I | EgAQP-SIP | Eucgr.B03790 | Eg | + | Chr02 | 57335757-57342026 | Nucleus |
| I | EgAQP-SIP | Eucgr.G02502 | Eg | + | Chr07 | 45835660-45839985 | Nucleus |
| II | AtAQP-NIP | AT1G31885 | At | + | Chr1 | 11450460-11451985 |  |
| II | AtAQP-NIP | AT1G80760 | At | - | Chr1 | 30350640-30352015 |  |
| II | AtAQP-NIP | AT2G29870 | At | + | Chr2 | 12741192-12741791 |  |
| II | AtAQP-NIP | AT2G34390 | At | - | Chr2 | 14514617-14515793 |  |
| II | AtAQP-NIP | AT3G06100 | At | - | Chr3 | 1841388-1842934 |  |
| II | AtAQP-NIP | AT4G10380 | At | - | Chr4 | 6431530-6434510 |  |
| II | AtAQP-NIP | AT4G18910 | At | + | Chr4 | 10366211-10368179 |  |
| II | AtAQP-NIP | AT4G19030 | At | - | Chr4 | 10421728-10423409 |  |
| II | AtAQP-NIP | AT5G37810 | At | + | Chr5 | 15045232-15047807 |  |
| II | AtAQP-NIP | AT5G37820 | At | + | Chr5 | 15050153-15051542 |  |
| II | EgAQP-NIP | Eucgr.D00421 | Eg | - | Chr04 | 7907818-7909769 | Cell membrane |
| II | EgAQP-NIP | Eucgr.D00422 | Eg | - | Chr04 | 7970919-7973125 | Cell membrane |
| II | EgAQP-NIP | Eucgr.D00423 | Eg | - | Chr04 | 7997249-7999767 | Cell membrane |
| II | EgAQP-NIP | Eucgr.D02548 | Eg | + | Chr04 | 39412291-39416681 | Cell membrane |
| II | EgAQP-NIP | Eucgr.E02324 | Eg | + | Chr05 | 35924878-35927744 | Cell membrane/Vacuole |
| II | EgAQP-NIP | Eucgr.E02325 | Eg | + | Chr05 | 35899998-35903001 | Cell membrane |
| II | EgAQP-NIP | Eucgr.E02326 | Eg | + | Chr05 | 35887332-35890323 | Cell membrane |
| II | EgAQP-NIP | Eucgr.E02327 | Eg | + | Chr05 | 35864700-35867725 | Cell membrane |
| II | EgAQP-NIP | Eucgr.E02616 | Eg | - | Chr05 | 40691671-40694468 | Cell membrane |
| II | EgAQP-NIP | Eucgr.E02836 | Eg | - | Chr05 | 47182717-47185380 | Cell membrane |
| II | EgAQP-NIP | Eucgr.E02837 | Eg | - | Chr05 | 47165072-47165900 | Cell membrane |
| II | EgAQP-NIP | Eucgr.E02838 | Eg | - | Chr05 | 47148225-47149417 | Cell membrane |
| II | EgAQP-NIP | Eucgr.F02826 | Eg | - | Chr06 | 40455259-40458174 | Cell membrane |
| II | EgAQP-NIP | Eucgr.G02188 | Eg | + | Chr07 | 41946068-41947932 | Cell membrane/Vacuole |
| II | EgAQP-NIP | Eucgr.H02230 | Eg | - | Chr08 | 28156882-28158912 | Cell membrane |
| II | EgAQP-NIP | Eucgr.H03586 | Eg | + | Chr08 | 49482102-49483300 | Cell membrane |
| II | EgAQP-NIP | Eucgr.J01345 | Eg | - | Chr10 | 15153835-15157905 | Cell membrane |
| II | EgAQP-NIP | Eucgr.J03103 | Eg | - | Chr10 | 36522801-36525292 | Cell membrane |
| III | AtAQP-TIP | AT1G17810 | At | + | Chr1 | 6130209-6131442 |  |
| III | AtAQP-TIP | AT1G52180 | At | - | Chr1 | 19424944-19425928 |  |
| III | AtAQP-TIP | AT1G73190 | At | + | Chr1 | 27522155-27523171 |  |
| III | AtAQP-TIP | AT2G25810 | At | + | Chr2 | 11012658-11013906 |  |
| III | AtAQP-TIP | AT2G36830 | At | + | Chr2 | 15445490-15446336 |  |
| III | AtAQP-TIP | AT3G16240 | At | + | Chr3 | 5505534-5506788 |  |
| III | AtAQP-TIP | AT3G26520 | At | - | Chr3 | 9722770-9723703 |  |
| III | AtAQP-TIP | AT3G47440 | At | + | Chr3 | 17482054-17482987 |  |
| III | AtAQP-TIP | AT4G01470 | At | - | Chr4 | 625092-625850 |  |
| III | AtAQP-TIP | AT4G17340 | At | + | Chr4 | 9699318-9700250 |  |
| III | AtAQP-TIP | AT5G47450 | At | - | Chr5 | 19248509-19249466 |  |
| III | EgAQP-TIP | Eucgr.B02403 | Eg | - | Chr02 | 42833697-42834827 | Vacuole |
| III | EgAQP-TIP | Eucgr.C02914 | Eg | + | Chr03 | 58649219-58650626 | Vacuole |
| III | EgAQP-TIP | Eucgr.D02090 | Eg | + | Chr04 | 34405445-34406458 | Vacuole |
| III | EgAQP-TIP | Eucgr.D02507 | Eg | - | Chr04 | 39054993-39056837 | Cell membrane/Vacuole |
| III | EgAQP-TIP | Eucgr.F03054 | Eg | + | Chr06 | 42512098-42513575 | Vacuole |
| III | EgAQP-TIP | Eucgr.H00668 | Eg | + | Chr08 | 9215306-9216722 | Vacuole |
| III | EgAQP-TIP | Eucgr.J00051 | Eg | - | Chr10 | 767801-769143 | Vacuole |
| III | EgAQP-TIP | Eucgr.J02074 | Eg | - | Chr10 | 25938193-25939503 | Vacuole |
| III | EgAQP-TIP | Eucgr.K00164 | Eg | + | Chr11 | 1075222-1076485 | Cell membrane/Vacuole |
| III | EgAQP-TIP | Eucgr.K02339 | Eg | + | Chr11 | 31191317-31193166 | Vacuole |
| III | EgAQP-TIP | Eucgr.K02438 | Eg | + | Chr11 | 32418501-32419341 | Vacuole |
| III | EgAQP-TIP | Eucgr.L03165 | Eg | + | scaffold_2175 | 2478-3874 | Vacuole |
| III | EgAQP-TIP | Eucgr.L03254 | Eg | - | scaffold_2473 | 198-2053 | Vacuole |
| III | EgAQP-TIP | Eucgr.L03444 | Eg | + | scaffold_3187 | 48-1181 | Vacuole |
| III | EgAQP-TIP | Eucgr.L00814 | Eg | - | scaffold_56 | 352041-354716 | Cell membrane |
| IV | AtAQP-PIP | AT1G01620 | At | - | Chr1 | 225986-227176 |  |
| IV | AtAQP-PIP | AT2G16850 | At | + | Chr2 | 7301647-7303180 |  |
| IV | AtAQP-PIP | AT2G37170 | At | - | Chr2 | 15613624-15614920 |  |
| IV | AtAQP-PIP | AT2G37180 | At | + | Chr2 | 15617779-15618937 |  |
| IV | AtAQP-PIP | AT2G39010 | At | + | Chr2 | 16291564-16292740 |  |
| IV | AtAQP-PIP | AT2G45960 | At | + | Chr2 | 18910450-18911703 |  |
| IV | AtAQP-PIP | AT3G53420 | At | - | Chr3 | 19803906-19805454 |  |
| IV | AtAQP-PIP | AT3G54820 | At | + | Chr3 | 20302117-20303738 |  |
| IV | AtAQP-PIP | AT3G61430 | At | + | Chr3 | 22733657-22735113 |  |
| IV | AtAQP-PIP | AT4G00430 | At | - | Chr4 | 186143-187531 |  |
| IV | AtAQP-PIP | AT4G23400 | At | + | Chr4 | 12220792-12222155 |  |
| IV | AtAQP-PIP | AT4G35100 | At | + | Chr4 | 16708672-16709958 |  |
| IV | AtAQP-PIP | AT5G60660 | At | - | Chr5 | 24375673-24376939 |  |
| IV | EgAQP-PIP | Eucgr.A01153 | Eg | - | Chr01 | 24874608-24877259 | Cell membrane |
| IV | EgAQP-PIP | Eucgr.B02499 | Eg | - | Chr02 | 45012796-45014028 | Cell membrane |
| IV | EgAQP-PIP | Eucgr.B02500 | Eg | - | Chr02 | 45017683-45018847 | Cell membrane |
| IV | EgAQP-PIP | Eucgr.D01836 | Eg | + | Chr04 | 31870718-31873051 | Cell membrane |
| IV | EgAQP-PIP | Eucgr.E00628 | Eg | + | Chr05 | 6027646-6029802 | Cell membrane |
| IV | EgAQP-PIP | Eucgr.E03494 | Eg | + | Chr05 | 61687177-61688466 | Cell membrane |
| IV | EgAQP-PIP | Eucgr.G03037 | Eg | - | Chr07 | 51562224-51563760 | Cell membrane |
| IV | EgAQP-PIP | Eucgr.I01369 | Eg | - | Chr09 |  | Cell membrane |
| IV | EgAQP-PIP | Eucgr.I01429 | Eg | + | Chr09 | 24647541-24649041 | Cell membrane |
| IV | EgAQP-PIP | Eucgr.J00237 | Eg | + | Chr10 | 2447356-2449192 | Cell membrane |
| IV | EgAQP-PIP | Eucgr.J00930 | Eg | + | Chr10 | 10150845-10152248 | Cell membrane |
| IV | EgAQP-PIP | Eucgr.J01087 | Eg | + | Chr10 | 11877465-11880807 | Cell membrane |
| IV | EgAQP-PIP | Eucgr.K02993 | Eg | + | Chr11 | 37413395-37414731 | Cell membrane |
| IV | EgAQP-PIP | Eucgr.K03244 | Eg | - | Chr11 | 40541781-40543093 | Cell membrane |
| IV | EgAQP-PIP | Eucgr.L02985 | Eg | + | scaffold_1745 | 3908-5126 | Cell membrane |

### *EgAQP* Genes Features

The chromosomal map of the *EgAQPs* was drawn using location information of the *Eucalyptus grandis*. The 48 *EgAQP* genes were widely distributed on the eleven *Eucalyptus grandis* chromosomes and five *EgAQP* genes were distributed on four scaffolds (**Supplementary Figure 1**). Chromosome 5 harbored the most *EgAQP* genes (ten total), followed by chromosome 10 and chromosome 7 (seven genes each) with the remaining *EgAQP* genes distributed along chromosomes. The genomic analysis continued with the genes encoding proteins harboring the entire MIP domain. Here, we have identified 14 *EgAQPs* originating from tandem duplications (TDs) within the *Eucalyptus grandis* genome. These TDs encompass specific genetic regions, including Eucgr.A02066 and Eucgr.A02068 (members of the *EgAQP-SIP* subfamily) located on chromosome 1, Eucgr.B02499 and Eucgr.B02500 (a part of the *EgAQP-PIP* subfamily) found on chromosome 2, and Eucgr.D00421, Eucgr.D00422, and Eucgr.D00423 (members of the *EgAQP-NIP* subfamily) situated on chromosome 4. Additionally, chromosome 5 hosts a cluster of genes, namely Eucgr.E02324, Eucgr.E02325, Eucgr.E02326, Eucgr.E02327, Eucgr.E02836, Eucgr.E02837, and Eucgr.E02838, all belonging to the *EgAQP-NIP* subfamily (see **Table 2**).

**Table 2.** Summary of the synteny between *E. grandis* and *E. grandis x urophylla*. Ka/Ks ratios for pairwise comparisons among members of the EgAQP subfamilies in Eucalyptus genus. Duplication and Purification selection are presented.

| AQP subfamily | ID *Eucalyptus grandis* | ID *Eucalyptus urophylla* x *grandis* | Ks | Kn | Kn/Ks | Purification selection | Duplication event |
| --- | --- | --- | --- | --- | --- | --- | --- |
| EgAQP-PIP | Eucgr.A01153 | Eug01G0099700 | 0.0235 | 0.0000 | 0.0000 | yes |  |
| EgAQP-SIP | Eucgr.A02066 | Eug01G0182900 | 0.0051 | 0.0019 | 0.3725 | yes | Tandem |
| EgAQP-SIP | Eucgr.A02068 | Eug01G0183000 | 0.0000 | 0.0000 | 0.0000 | yes | Tandem |
| EgAQP-SIP | Eucgr.A02176 | Eug01G0192500 | 0.0090 | 0.0022 | 0.2444 | yes |  |
| EgAQP-TIP | Eucgr.B02403 |  |  |  |  | yes |  |
| EgAQP-PIP | Eucgr.B02499a | Eug02G0206300 | 0.0179 | 0.0015 | 0.0838 | yes | Tandem |
| EgAQP-PIP | Eucgr.B02500 |  |  |  |  | yes | Tandem |
| EgAQP-SIP | Eucgr.B03790 | Eug02G0313800 | 0.0319 | 0.0085 | 0.2665 | yes |  |
| EgAQP-TIP | Eucgr.C02914 | Eug03G0220300 | 0.0482 | 0.0038 | 0.0788 | yes |  |
| EgAQP-NIP | Eucgr.D00421a |  |  |  |  | yes | Tandem |
| EgAQP-NIP | Eucgr.D00422a |  |  |  |  | yes | Tandem |
| EgAQP-NIP | Eucgr.D00423a | Eug04G0038100 | 0.2667 | 0.0661 | 0.2478 | yes | Tandem |
| EgAQP-PIP | Eucgr.D01836 | Eug04G0145500 | 0.0000 | 0.0000 | 0.0000 | yes |  |
| EgAQP-TIP | Eucgr.D02090 | Eug04G0166500 | 0.4232 | 0.1573 | 0.3717 | yes |  |
| EgAQP-TIP | Eucgr.D02507 | Eug04G0203700 | 0.0208 | 0.0138 | 0.6635 | yes |  |
| EgAQP-NIP | Eucgr.D02548 | Eug04G0206900 | 0.0083 | 0.0000 | 0.0000 | yes |  |
| EgAQP-PIP | Eucgr.E00628 | Eug05G0056800 | 0.1228 | 0.0686 | 0.5586 | yes |  |
| EgAQP-NIP | Eucgr.E02324a |  |  |  |  | yes | Tandem |
| EgAQP-NIP | Eucgr.E02325a |  |  |  |  | yes | Tandem |
| EgAQP-NIP | Eucgr.E02326a |  |  |  |  | yes | Tandem |
| EgAQP-NIP | Eucgr.E02327a | Eug05G0174200 | 0.0086 | 0.0178 | 2.0698 | no | Tandem |
| EgAQP-NIP | Eucgr.E02616 | Eug05G0174200 | 0.0449 | 0.0237 | 0.5278 | yes |  |
| EgAQP-NIP | Eucgr.E02836a | Eug05G0184900 | 0.1495 | 0.0347 | 0.2321 | yes | Tandem |
| EgAQP-NIP | Eucgr.E02837a |  |  |  |  | yes | Tandem |
| EgAQP-NIP | Eucgr.E02838a |  |  |  |  | yes | Tandem |
| EgAQP-PIP | Eucgr.E03494 | Eug05G0226800 | 0.2139 | 0.1184 | 0.5535 | yes |  |
| EgAQP-NIP | Eucgr.F02826 | Eug06G0239400 | 0.8973 | 0.6580 | 0.7333 | yes |  |
| EgAQP-TIP | Eucgr.F03054 | Eug06G0261400 | 0.0040 | 0.0000 | 0.0000 | yes |  |
| EgAQP-NIP | Eucgr.G02188 | Eug07G0185700 | 0.0474 | 0.0272 | 0.5738 | yes |  |
| EgAQP-SIP | Eucgr.G02502 | Eug07G0212700 | 0.0061 | 0.0033 | 0.5410 | yes |  |
| EgAQP-PIP | Eucgr.G03037 | Eug07G0260400 | 0.0303 | 0.0000 | 0.0000 | yes |  |
| EgAQP-TIP | Eucgr.H00668 | Eug08G0060900 | 0.0621 | 0.0035 | 0.0564 | yes |  |
| EgAQP-NIP | Eucgr.H02230 |  |  |  |  | yes |  |
| EgAQP-NIP | Eucgr.H03586 | Eug08G0250500 | 0.1250 | 0.0327 | 0.2616 | yes |  |
| EgAQP-NIP | Eucgr.I01369 | Eug09G0117900 | 0.0099 | 0.0027 | 0.2727 | yes |  |
| EgAQP-PIP | Eucgr.I01429 | Eug09G0123800 | 0.0484 | 0.0016 | 0.0331 | yes |  |
| EgAQP-TIP | Eucgr.J00051 | Eug10G0005500 | 0.0292 | 0.0000 | 0.0000 | yes |  |
| EgAQP-PIP | Eucgr.J00237 | Eug10G0022900 | 0.0089 | 0.0102 | 1.1461 | no |  |
| EgAQP-PIP | Eucgr.J00930 | Eug10G0084700 | 0.0119 | 0.0090 | 0.7563 | yes |  |
| EgAQP-PIP | Eucgr.J01087 | Eug10G0100000 | 0.0176 | 0.0016 | 0.0909 | yes |  |
| EgAQP-NIP | Eucgr.J01345 | Eug10G0122900 | 0.0054 | 0.0023 | 0.4259 | yes |  |
| EgAQP-TIP | Eucgr.J02074 | Eug10G0196000 | 0.0000 | 0.0031 | 0.0000 | yes |  |
| EgAQP-NIP | Eucgr.J03103 | Eug10G0279100 | 563685 | 12933 | 0.0229 | yes |  |
| EgAQP-TIP | Eucgr.K00164 | Eug11G0009300 | 0.0128 | 0.0019 | 0.1484 | yes |  |
| EgAQP-TIP | Eucgr.K02339 | Eug11G0185100 | 0.0562 | 0.0018 | 0.0320 | yes |  |
| EgAQP-TIP | Eucgr.K02438 | Eug11G0190600 | 0.0455 | 0.0000 | 0.0000 | yes |  |
| EgAQP-PIP | Eucgr.K02993 | Eug11G0235800 | 0.0099 | 0.0000 | 0.0000 | yes |  |
| EgAQP-PIP | Eucgr.K03244 | Eug11G0254300 | 0.0338 | 0.0013 | 0.0385 | yes |  |

Using the Plaza database, our analysis has unveiled that TDs on chromosomes 1 and 2 are shared between *Arabidopsis thaliana* and *Eucalyptus grandis*, signifying their evolutionary conservation. Notably, *EgAQP-NIP* exhibits a notably higher TD ratio compared to the other *EgAQPs*. To investigate these phenomena within the Eucalyptus genus, we conducted a synteny analysis and calculated the synonymous (Ks) and nonsynonymous (Kn) substitution rates, resulting in the Kn/Ks ratios, for 53 *EgAQP* genes through pairwise comparisons between *Eucalyptus grandis* and *Eucalyptus urophylla* x *grandis* (as detailed in **Table 2**). Our findings indicate that the TDs of *EgAQP-NIPs* are unique to the *Eucalyptus grandis* genome, as outlined in **Table 2**. Furthermore, a limited number of genes exhibit evidence of positive selection (Kn/Ks > 1), exemplified by Eucgr.E02327 (Kn/Ks = 2.07) and Eucgr.J00237 (Kn/Ks = 1.15), while the remaining genes appear to undergo purified selection (**Table 2**).

### EgAQP Protein and Motifs Analysis

A multiple sequence alignment of the protein sequences revealed the existence of a functional MIP domain (IPR000425), with limited conservation across the EgAQPs (**Figure 2**), maintaining only the characteristic Asn-Pro motifs starting at position 272 and 479, both sites of NPA box (Froger *et al*. 1998) even when the multiple alignment is performed isolated for each subfamily (**Supplementary Figures 2-5**). Our investigation into the diversification of the protein sequence of the EgAQP subfamilies encompassed the analysis of motifs within the 53 identified proteins. However, two proteins (Eucg.A02068 and Eucg.B03790) lack conserved motifs compared with all EgAQP-SIP sets. Remarkably, the composition of these motifs exhibited significant diversity among the EgAQP subfamilies, resulting in the identification of a total of six conserved motifs (motif-1, motif-2, motif-3, motif-4, motif-5, and motif-7) associated with the MIP domain (**Figure 3A**). While these motifs are generally present in all EgAQPs, two additional motifs (motif-8 and motif-12) are unique to the AQP-NIP subfamily, and three other motifs (motif-9, motif-10, and motif-11) have been exclusively associated with the AQP-PIP subfamily (**Figure 3**). Furthermore, our analyses indicate a discernible pattern of motif diversification and reorganization within individual EgAQP subfamilies resulting in the variation of N-terminal, C-terminal, and domain protein (**Figure 3B**).

**Figure 2.**
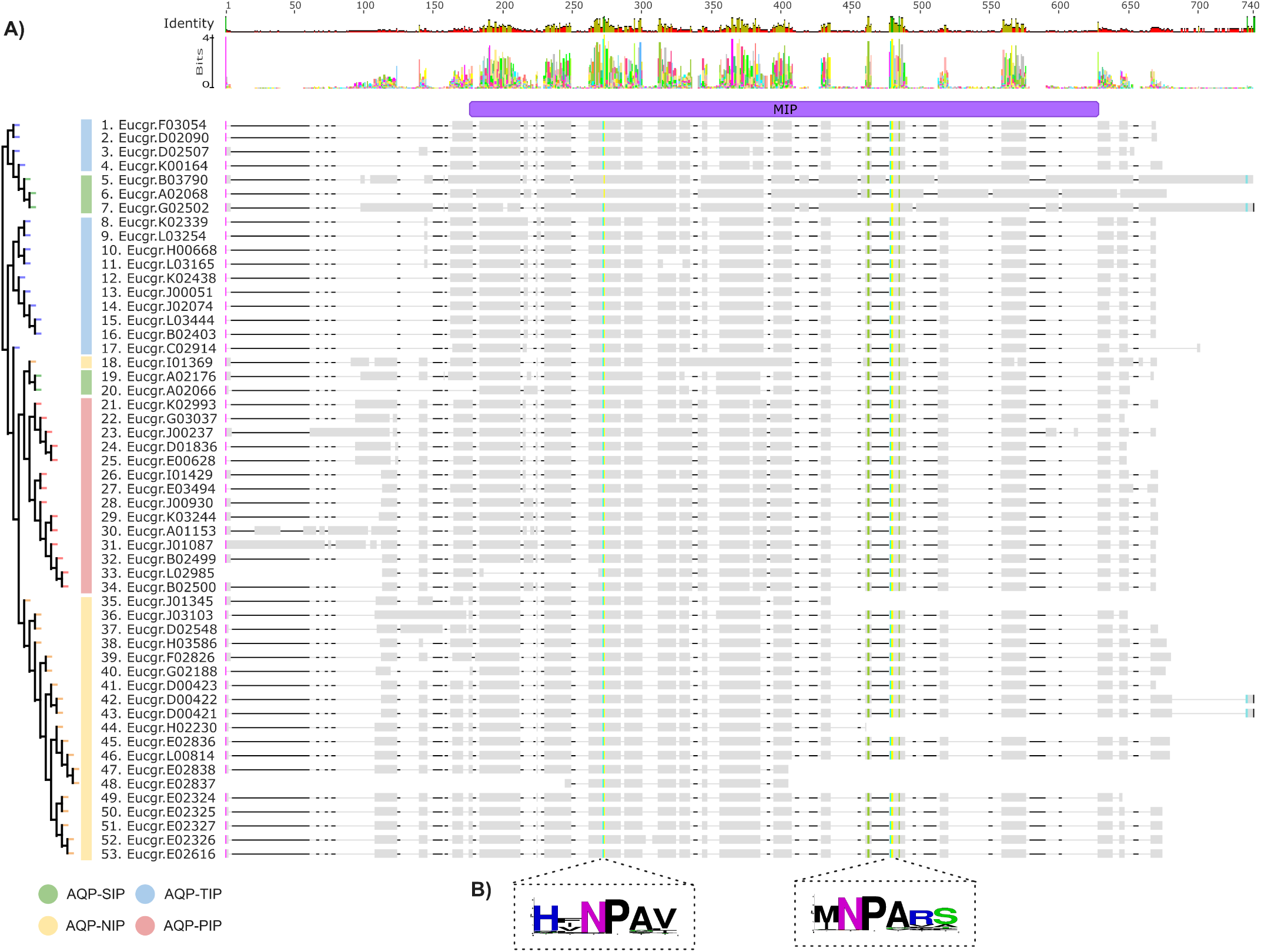
The sequences of AQPs from *Eucalyptus grandis* underwent a global multiple alignment constructed using MAFFT v7. This alignment highlights the annotation of the Major Intrinsic Protein (MIP) domain across all proteins and low conservation degrees along protein homologs. It also underscores the conservation of the NPA-box residues within the AQPs.

**Figure 3.**
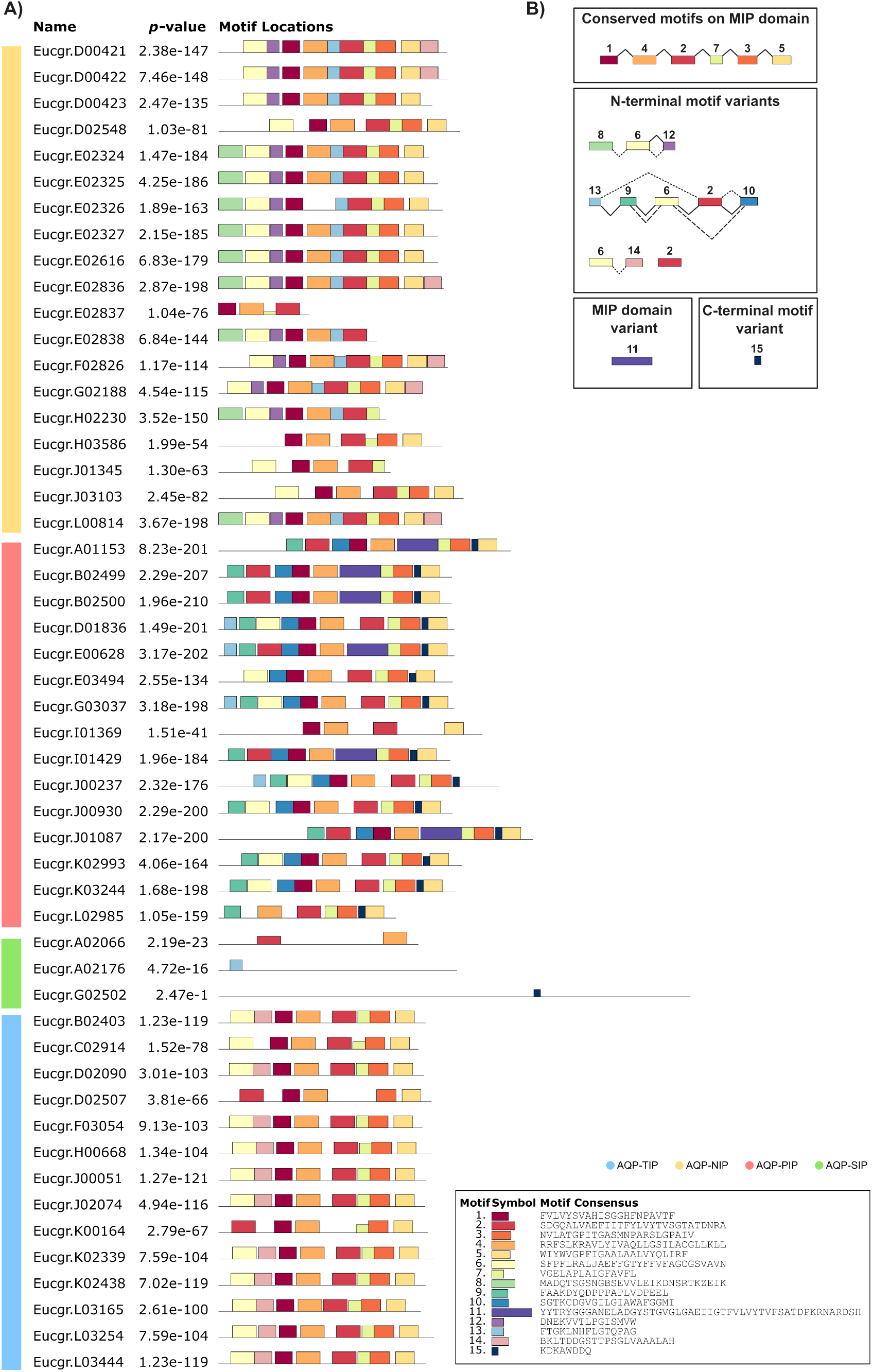
The conserved motifs were analyzed in EgAQPs using the MEME suite, with parameters set to extract 15 distinct conserved motifs. **A)** These motifs are depicted with their respective consensus sequences, alongside the AQP subfamilies identified in the left sidebar, as indicated by the subtitle. **B)** Conserved and variation motifs sets found in EgAQP proteins.

### EgAQPs Phylogenetic Relationships Analysis

To elucidate the phylogeny among EgAQP proteins, the aLRT SHlike algorithm was employed to construct a phylogenetic tree encompassing all AQP sequences with the entire MIP domain from both *Eucalyptus grandis* and *Arabidopsis thaliana* (**Figure 1**). The outcome of the phylogenetic analysis revealed a putative plant AQP phylogeny, congruent with eudicot species. This observation implies a relatively recent divergence from a common ancestor to eudicots compared to the lineage of non-seed plants (*Physcomitrium patens* as outgroup). Notably, four distinct AQP clades (clade I-SIP, clade II-NIP, clade III-TIP, and clade IV-PIP) were discerned, implying that the expansion of specific AQP gene subfamilies might have been facilitated by whole-genome duplication or segmental duplications (**Figure 1** and **Table 2**). Additionally, the variable distribution of *EgAQP* members across subfamilies, as observed in the phylogenetic analysis (**Figure 1**) and summarized in Table 1, demonstrates that *EgAQP-SIP* comprises 5 members, *EgAQP-NIP* consists of 19 members, *EgAQP-TIP* encompasses 14 members, and *EgAQP-PIP* encompasses 14 members.

### Expression Profile of *AQP*s in *Arabidopsis thaliana* and *Eucalyptus grandis* Across Tissues

To identify and profile the expression of *AQP*s, the transcript levels in the root, leaf, stem, carpel, pollen, aerial part, receptacle, and the gynoecium of *Arabidopsis thaliana*, along with transcripts for leaf, stem, and root in *Eucalyptus grandis* were accessed. A heatmap comparing the *AQP* expression patterns within and between samples is presented in **Figure 5** (expression level in log(TPM+1)). Analysis of the expression of *AQP*s in different tissues/organs revealed a higher expression of *SIP* and *PIP* followed by *TIP* and *NIP* across all tissues examined in both species. A notable observation is *PIP* genes’ relatively higher expression than *TIP* genes, mostly in vascular tissues such as roots and stems. Certain genes, such as Eucgr.J00930, Eucgr.D01836, and Eucgr.E00628 stand out with pronounced expression levels (**Figure 5**). At the same time, the *TIP* and *NIP* groups have genes such as Eucgr.L03254 (TIP), Eucgr.L03165 (TIP), Eucgr.H00668 (TIP), Eucgr.H03586 (NIP), Eucgr.g02188 (NIP) and Eucgr.F02826 (NIP) with no expression in *Eucalyptus grandis*, the same is observed in *Arabidopsis thaliana* orthologs. Conversely, the orthologs demonstrate high expression across the root, stem, and leaf tissues. For instance, the genes AT3G53420 (*AtPIP2;1*), AT2G25810 (*AtTIP4;1*), and AT1G01620 (*AtPIP1;3*), and their corresponding orthologs Eucgr.A01153, Eucgr.K03244, Eucgr.C02914, Eucgr.D01836, Eucgr.E00628, and Eucgr.K02993, exhibit high expression in the three least vascularized tissues (leaves, stem, and roots). On the other hand, for the gene AT4G35100 (*AtPIP3*), its orthologs Eucgr.I01429 and Eucgr.J00930 also show a constant expression pattern across root, leaf, and stem tissues. Certain genes such as the *EgAQP-PIP* in *Eucalyptus grandis* display constitutive expression profiles in all analyzed tissues and are likely housekeeping genes necessary for basic metabolic functions across tissues.

### Expression Profile of *EgAQP*s Under Drought Stress and Rewatering

To uncover potential homologs of *AQP* genes associated with responses to drought stress, we conducted an analysis of the RNAseq dataset for *Eucalyptus grandis* exposed to drought stress (DS) and subsequent rewatering (RW) conditions as described in Teshome *et al*. (2023). We employed a significance threshold with an adjusted p-value of less than 0.05 and a log fold change (logFC) greater than 2 to identify differentially expressed genes (DEGs). The drought stress resulted in alterations to the expression profiles of *EgAPQs* in *Eucalyptus grandis*. Specifically, we observed up-regulation in the expression profiles of *EgAQP-TIP* (Eucgr.C02914) and *EgAQP-NIP* (Eucgr.D02548). Conversely, three members of *EgAQP-PIP* (Eucgr.A01153, Eucgr.G03037, and Eucgr.K03244) displayed down-regulation during the drought stress phase. However, when the plants were rewatered, these same genes exhibited an opposite expression profile, including an additional member, *EgAQP-PIP* (Eucgr.E00628), which was up-regulated (refer to **Figure 6A**). We delved into numerous studies concerning AQPs and their potential involvement in water transport across various subcellular compartments, as depicted in **Figure 6B**. Furthermore, we utilized Plant-mPloc (Chou and Shen 2010) to predict the subcellular localization of six differentially expressed *EgAQPs* (**Table 1**). Our findings suggest that *EgAQP-TIP* (Eucgr.C02914) may be targeted to the vacuole, while *EgAQP-NIP* (Eucgr.D02548) and *EgAQP-PIP* (Eucgr.A01153, Eucgr.G03037, and Eucgr.K03244) may be associated with the plasma membrane. These AQP subfamilies have been previously linked to cellular water transport, as shown in **Figure 6B**.

## DISCUSSION

### Genomic Abundance and Diversification of Aquaporin Genes in Plants

Numerous species, whose *AQP*s have been characterized at the genomic level, possess a relatively high number of those *AQP* genes. In the case of *Eucalyptus grandis*, our study identified and characterized 53 genes belonging to the *AQP* family. As the AQPs are largely a protein superfamily, (Ishibashi *et al*. 2020) we classified the *EgAQPs* into four main subfamilies (AQP-SIP, AQP-NIP, AQP-TIP, and AQP-PIP) which have resulted due to increasing expansion and diversification through successive rounds of duplication of the genome.

In addition to *Eucalyptus grandis*, several plant species exhibit a notable abundance of *AQP* genes. In the case of *Brassica napus*, a remarkable 121 *AQP* genes have been reported (Yuan *et al*. 2017). Conversely, *Arabidopsis thaliana*, which also belongs to the Brassicaceae family, possesses only 35 *AQP* isoforms distributed across five chromosomes (Johanson *et al*. 2001). The genus *Gossypium* sp. has a substantial presence of 71 *AQPs* (Li *et al*. 2019), whereas *Acacia auriculiformis* has a more modest count of 21 *AQPs* (Zhang *et al*. 2021). Another hand, *Vitis vinifera* L. contains 28 *AQP* genes Fouquet *et al*. (2008). In the order Malpighiales, species such as *Populus trichocarpa* contain 55 *AQP* homologs (Gupta and Sankararamakrishnan 2009), while *Manihot esculenta* possesses 45 (Putpeerawit *et al*. 2017). In monocots, *Oryza sativa* with 33 *AQP* genes (Sakurai *et al*. 2005). The greater abundance of AQP genes implies an association with the whole-genome duplication (WGD) ratio, as has been reported in the cases of *Brassica* and *Gossypium* genus (see the review by Liang and Schnable (2018)).

The variation in the count of *AQP* genes among vascular plants can be attributed to TD and WGD events, which are likely responsible for shaping the expansion and rearrangement of gene families in these organisms (Laloux *et al*. 2018). Moreover, angiosperms frequently exhibit tandem duplications that can further contribute to genomic rearrangements and expansions (Cannon *et al*. 2004; Zhang and Wang 2005). The sequencing of *Eucalyptus grandis* revealed that its genome contains the highest number of genes in tandem repeats when compared to all available sequenced plant genomes (Myburg *et al*. 2014). This phenomenon explains the elevated rates of tandem duplication observed in *EgAQPs*, particularly in the subfamilies *EgAPQ-SIP*, *EgAPQ-NIP*, and notably in *EgAPQ-PIP*.

### Functional Diversification of *EgAQPs*

Recent studies have identified two primary phylogenetic branches within AQPs, which include water-selective type channels (AQPs) and aquaglyceroproteins (GLPs) (Heymann and Engel 1999; Tong *et al*. 2019). This classification is based on distinctive amino acid sequences that show limited conservation around the NPA (Asn-Pro-Ala) sites, which are crucial for determining both large and small pore sizes. This division suggests an evolutionary origin rooted in Prokaryotes (Ishibashi *et al*. 2020; Bezerra-Neto *et al*. 2019). In Eukaryotes, both WGDs and TDs account for the majority of gene duplicates, leading to a significant number of paralogs in genomes undergoing diversification (Fang *et al*. 2012; HUANG *et al*. 2022). However, all of these new genes are not equal copies of the original. Most duplicated genes become neo-functionalized where they adopt a new function, while others become non-functional pseudogenes, and still more can be lost from the genome entirely (Sandve *et al*. 2018). We observed this trend in *Eucalyptus grandis* as only 48 of the 53 EgAQPs in the ortho family accession number ORTHO05D001001 contained a complete MIP domain, suggesting the presence of five pseudogenes (Figures 3-4, and Table 2).

**Figure 4.**
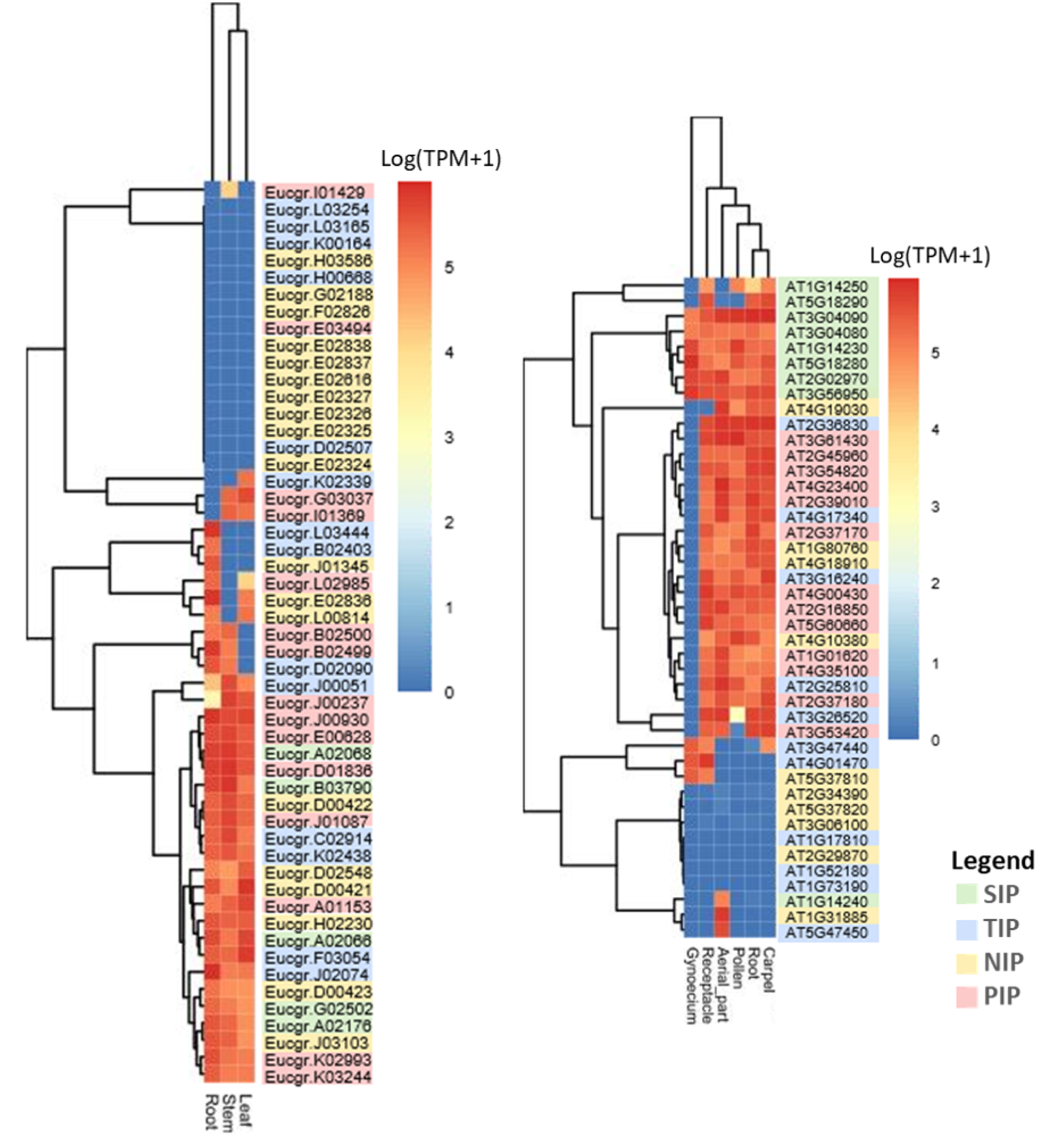
The expression patterns of EgAQPs and AtAQPs in various tissues through RNA-seq analysis are represented in log(TPM+1) values. On the left, we highlight the expression levels of *EgAQP* genes in leaf, stem, and root. On the right, we delve into the expression patterns of *AtAQP* genes in carpel, root, pollen, aerial part, receptacle, and gynoecium.

**Figure 5.**
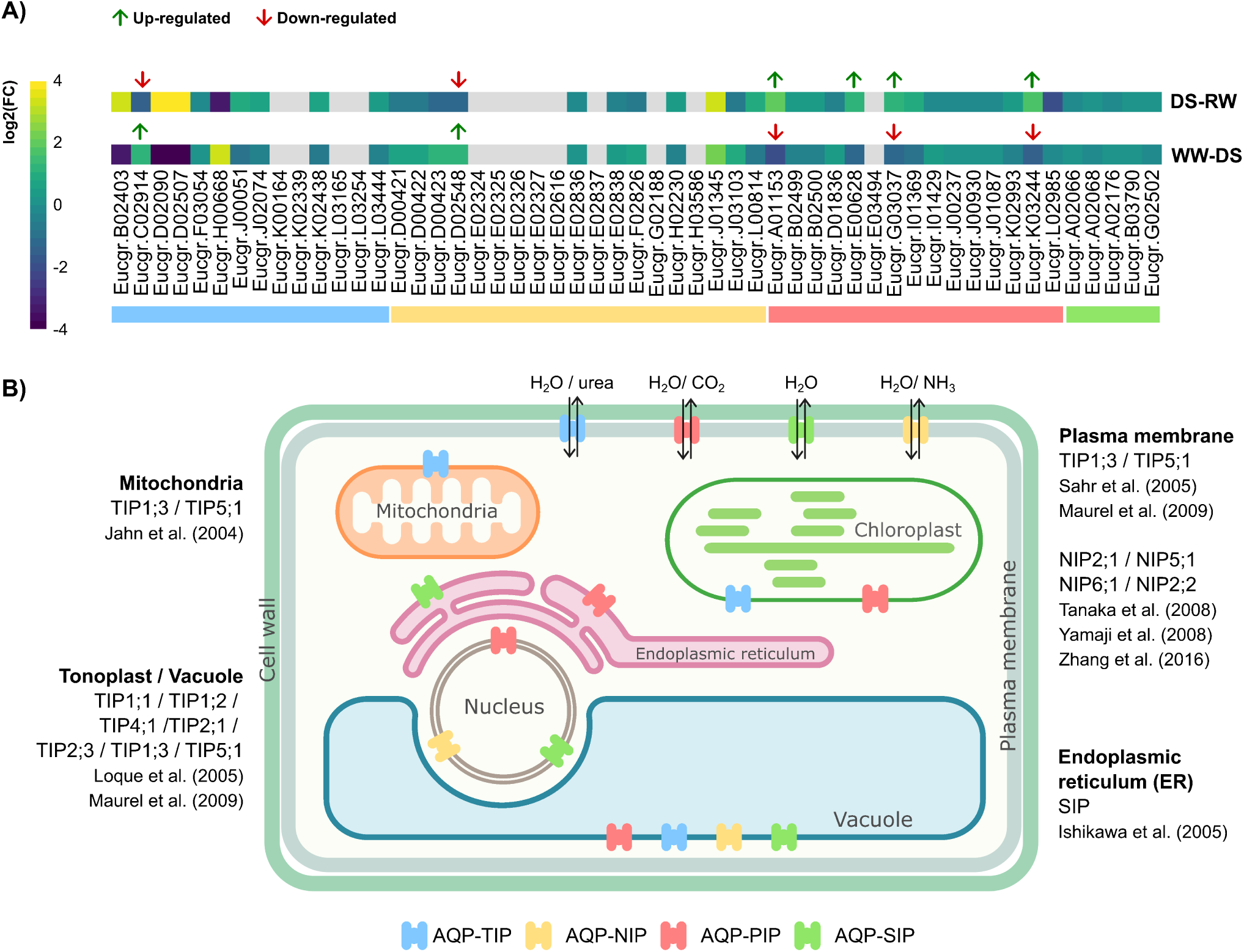
Schematic representation depicting the cellular localization of AQPs in a plant cell, as outlined in the literature. A) A heatmap showing the expression profile (logFC) of *EgAQPs* in an experiment of drought stress followed by the rewatering of *Eucalyptus grandis*. AQP genes overexpressed were represented by the arrows in their respective labels. Green arrows represent up-regulated genes while red arrows represent the down-regulated ones. **B)** AQPs are illustrated based on their subfamilies, indicating their presence within inner organelles and the plasma membrane based to literature.

Together with the phylogenetic analysis of the EgAQPs, along with their respective protein motifs, we have unveiled the expansion of AQP genes that are the result of significant differences as the species developed over evolutionary time (see the review by (Groszmann *et al*. 2017)). With this analysis, we can generate stronger hypotheses of how and why the EqAQPs are so diverse. The variation in functions and the emergence of new motifs is likely attributed to a series of independent gene duplication events along with the accumulation of asymmetric mutations (Sung *et al*. 2015). In addition to duplication events, we also show that AQP genes are subject to substantial mutations and can neofunctionalize due to selective pressure (**Figure 3**, **Table 2**). In most EgAQPs, the MIP domain (IPR000425) retains structural elements and motifs (1-5, and 7) (**Figure 3B**). However, many EQP-NIPs have obtained variations of an N-terminal targeting sequence (Figure 3B). Furthermore, some EgAQP-SIP sequences do not possess conserved AQP motifs and the motifs they do have are not shared across the subfamily as indicated by our parameter analysis (using an E-value threshold of *≤* 0.05). These unique characteristics of the EgAQP-SIPs have been previously reported, which supports our findings about this subfamily (Quigley *et al*. 2001).

Due to their diverse properties, AQP genes play crucial roles in plant growth, development, and stress responses (Afzal *et al*. 2016). Therefore, there has been significant interest in investigating the expression patterns of AQPs across different plant tissues. Here, we report a collective analysis of previous datasets on *AQP* expression in *Eucalyptus grandis* leaf, stem, and root tissue, along with carpel, root, pollen, aerial, gynoecium, and receptacle tissues in *Arabidopsis thaliana*. Our results show varying expression profiles among the *AQP* gene subfamilies. Both *Arabidopsis thaliana* and *Eucalyptus grandis* had high APQ expression in roots, however, no distinct patterns were identified between the two families. Consistent with *Arabidopsis thaliana*, many of the *EgAQP-NIP* and *EgAQP-TIP* genes had little to no detectable expression in the tissues analyzed in this study. This could be due to the specialized roles of *AQP-NIP*s in another development, pollen elongation, and seed germination, whose corresponding tissues were not measured here (Biela *et al*. 1999; Uehleln *et al*. 2003; Lopez *et al*. 2003; Bots *et al*. 2005; Durbak *et al*. 2014; Zhang *et al*. 2022). The TIP subfamily appears associated with various cellular signals, molecules such as urea and ammonia (Bezerra-Neto *et al*. 2019), additionally, these genes demonstrate distinct responses to water stress, salt, and cold conditions, as reported *Arabidopsis thaliana* (Afzal *et al*. 2016).

As for the AQP-SIP, even without having many extensive annotations, some have been functionally characterized. In *Arabidopsis thaliana*, an AQP-SIP has been linked to pollen germination and pollen tube elongation (Sato and Maeshima 2019). While our findings reveal a distinct expression pattern of AQP-SIPs in the gynoecium of *Arabidopsis thaliana*, a lack of publically available datasets for *Eucalyptus grandis* meant we could not investigate a potential shared expression pattern. The expression of *AQP-PIP* genes, however, remained consistent between both species and across various tissue types. This pattern of expression reinforces the existing literature showing AQP-PIPs are critical for maintaining and regulating water trafficking within cells Guo *et al*. (2006), Liu *et al*. (2020). The widespread expression patterns, subcellular targeting mechanisms, and the divergence from the functional domain collectively emphasize the high non-synonymous mutation rate of EgAQP, driving their diversification in *Eucalyptus grandis*.

### Drought Stress and Re-watering Affect the Abundance of EgAQP Transcripts

As the climate continues to change, limited water availability and high temperatures have resulted in drought stresses that have already begun to hinder the growth and productivity of the *Eucalyptus grandis* trees (Saharan *et al*. 2022). Here, we have identified six *EgAQPs* that play a pivotal role in responding to drought stress and subsequent rehydration (**Figure 6**. Additionally, we have shown an increase in the expression of *EgAQP-NIP* (Eucgr.D02548) and *EgAQP-TIP* (Eucgr.B02914) genes following 22 days of severe drought stress. This is in line with work in *A. thaliana*, which reports an increase in *AtAQP-TIP1;1* and *AtAQP-TIP2;1* transcript levels within 24 hours of drought (Feng *et al*. 2018). Both *AtAQP-TIP1;1* and *AtAQP-TIP2;1* encode proteins that facilitate water transport across the tonoplast to maintain the osmotic potential between the cytosol and vacuole (Rodrigues *et al*. 2013, 2016). Furthermore, the TIPs are known to enhance efficient transcellular water transport through live cells as needed (Barrieu *et al*. 1998). In contrast, *EgAQP-PIPs* (Eucgr.A01153, Eucgr.G03037, and Eucgr.K032) were down-regulated following the drought stress. This could result in reduced water permeability for the cell membranes as the transcripts code for proteins targeted to the cell membrane, which could either help to maintain cytoplasmic water levels or hinder the ability of water to enter the cell to prevent drought-induced dehydration and organellar hemifusion (Yaneff *et al*. 2014).

## CONCLUDING REMARKS

Here we identify and broadly characterize *EgAQPs* a set of 48 AQPs in *Eucalyptus grandis*. TD events were therefore the main driver of *EgAQP* diversity and expansion, specific to *Eucalyptus grandis* and not shared with *Eucalyptus grandis x urophylla*. Specifically, we have highlighted the *EgAQP-TIP*, *EgAQP-NIP*, and *EgAQP-PIP* isoforms, which showed a significant increase in expression during drought and subsequent re-watering. Further investigations are required to determine the precise relationship between specific AQPs and any potential roles in enhancing drought tolerance. In light of the pressing climatic crisis and changing environmental conditions worldwide, the cultivation of drought-tolerant plant varieties has become a paramount challenge in modern agriculture. The genes we have identified can serve as valuable resources for targeted engineering, thus paving the way for developing drought-resistant woody plants.

### Data availability

The datasets presented in this study can be found in online repositories SRA-NCBI (https://www.ncbi.nlm.nih.gov/ sra). The accession numbers are: PRJNA194429, PRJNA242915, PRJNA195608, PRJNA151589, PRJNA231089, PRJNA234023, and PRJNA896601.

## Acknowledgments

We would like to express our heartfelt gratitude to Zachery and Ravi for their invaluable contributions to this manuscript. Their unwavering support, guidance, and dedication have been instrumental in the successful completion of this work. In the pursuit of scientific knowledge, this endeavor has been greatly enriched by the principles of open science. We extend our appreciation to the global scientific community for promoting transparency, collaboration, and the free exchange of ideas. This open approach has been the cornerstone of our research, allowing us to build upon the collective wisdom of experts worldwide. Moreover, we are indebted to the advocates of public availability of data, as their efforts have provided us with a robust foundation for our research. The wealth of publicly accessible data sources has enabled us to explore new frontiers in our scientific exploration, fostering innovation and discovery. We recognize and acknowledge that this work could not have been accomplished without the collective support, inspiration, and resources that the field of science has to offer. It is with profound gratitude that we acknowledge the interconnected efforts of researchers, institutions, and individuals who have championed open science and public data availability.

## Author’s contributions

DSS, PHC, and GS analyzed the data, prepared figures and/or tables, authored or reviewed article drafts, and approved the final draft. SNW reviewed drafts of the article and approved the final draft. RVM and ZDS conducted additional analyses, reviewed article drafts, and approved the final draft. HMD conceived and designed the workflow of analyses data, prepared figures and/or tables, authored or reviewed article drafts, and approved the final draft.

---

**Figure S1.**
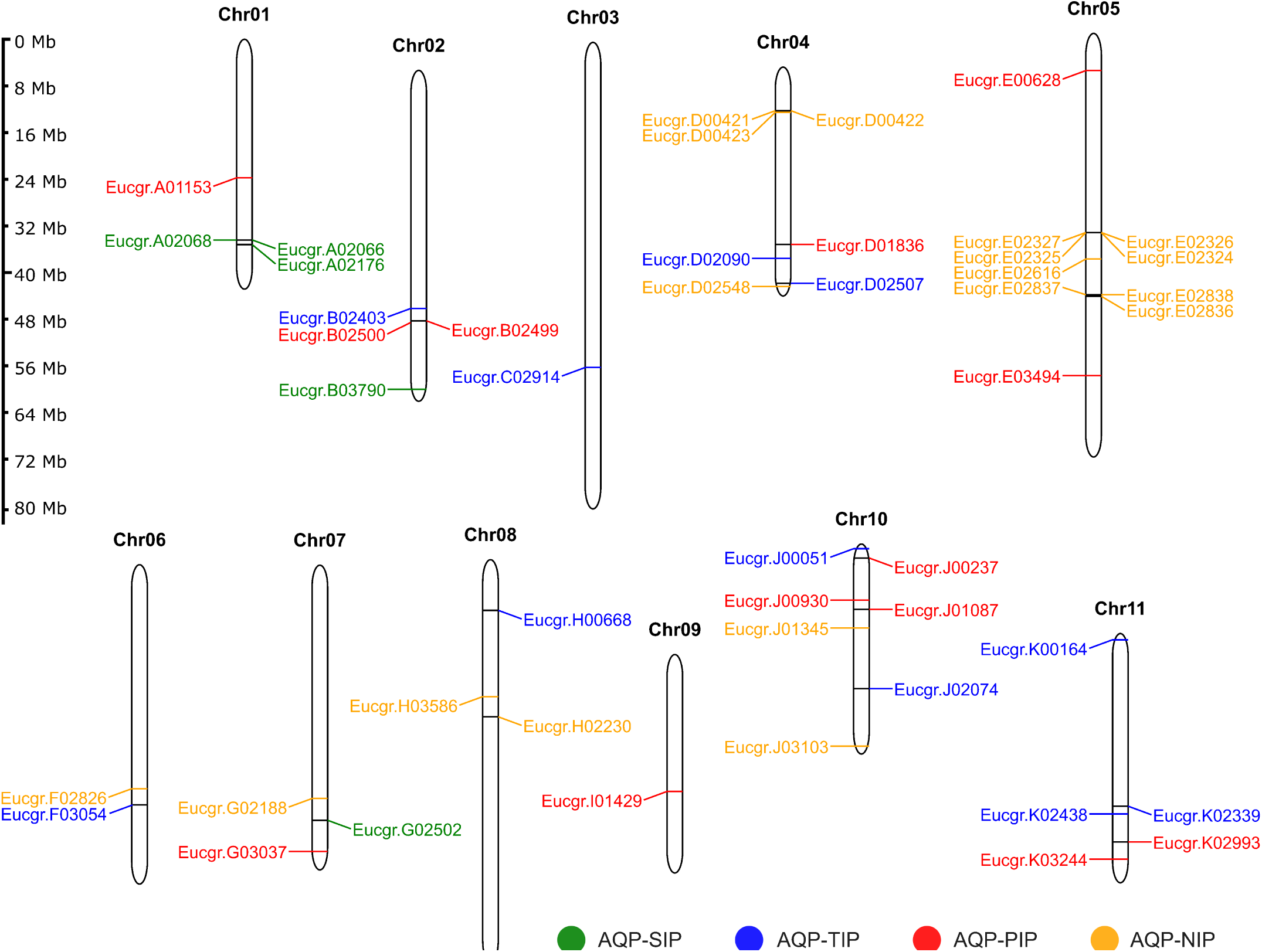
Chromosomal distribution of aquaporin genes in *Eucalyptus grandis*. The four protein subfamilies received different color: Green is for SIP, blue for TIP, NIP is yellow and PIP is red. The side bar depicts the chromosome size in a scale of megabases. The abbreviations used are Ch (chromosome) and Mb (megabase).

**Figure S2.**
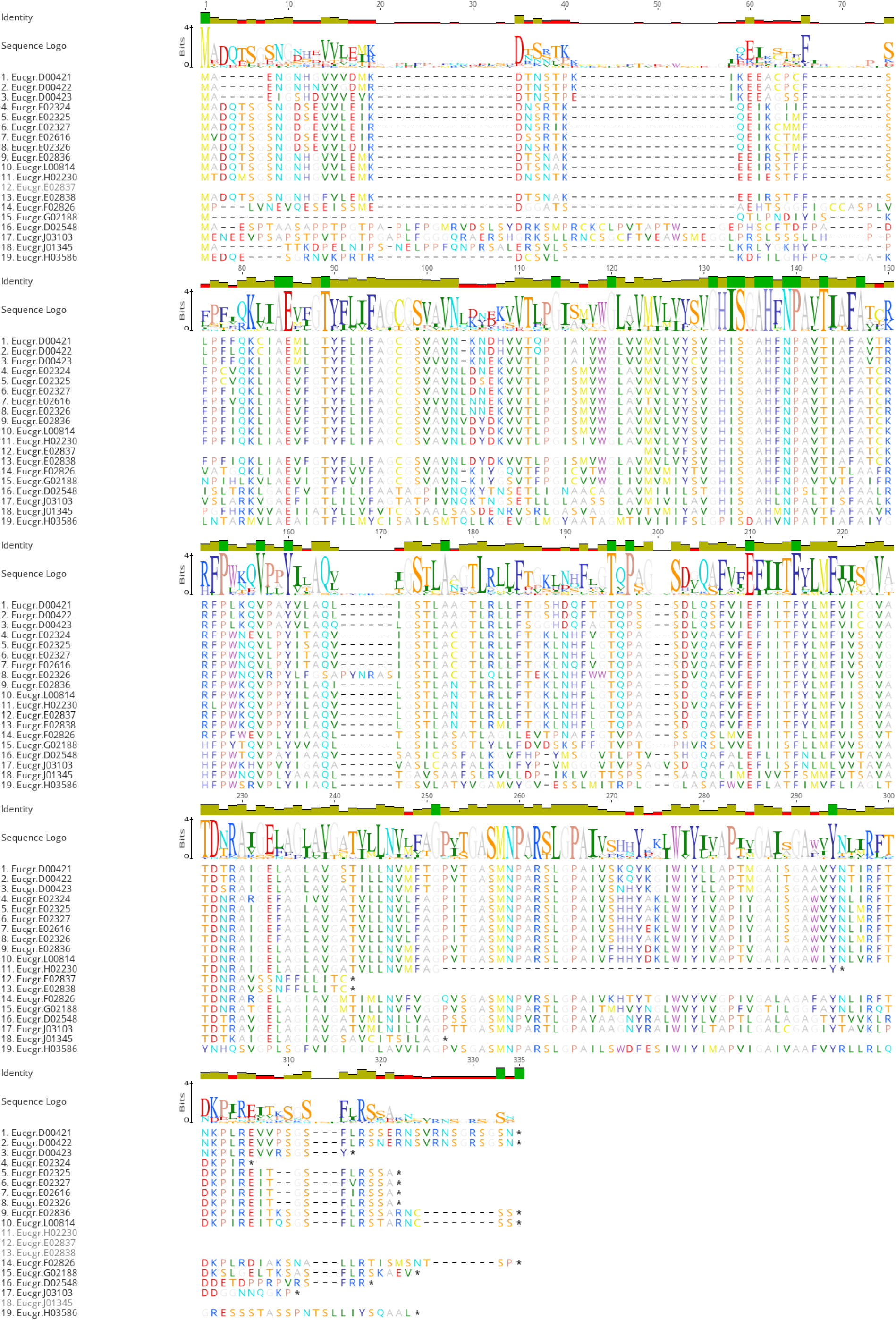
Multiple alignment of EgAQP-NIP proteins.

**Figure S3.**
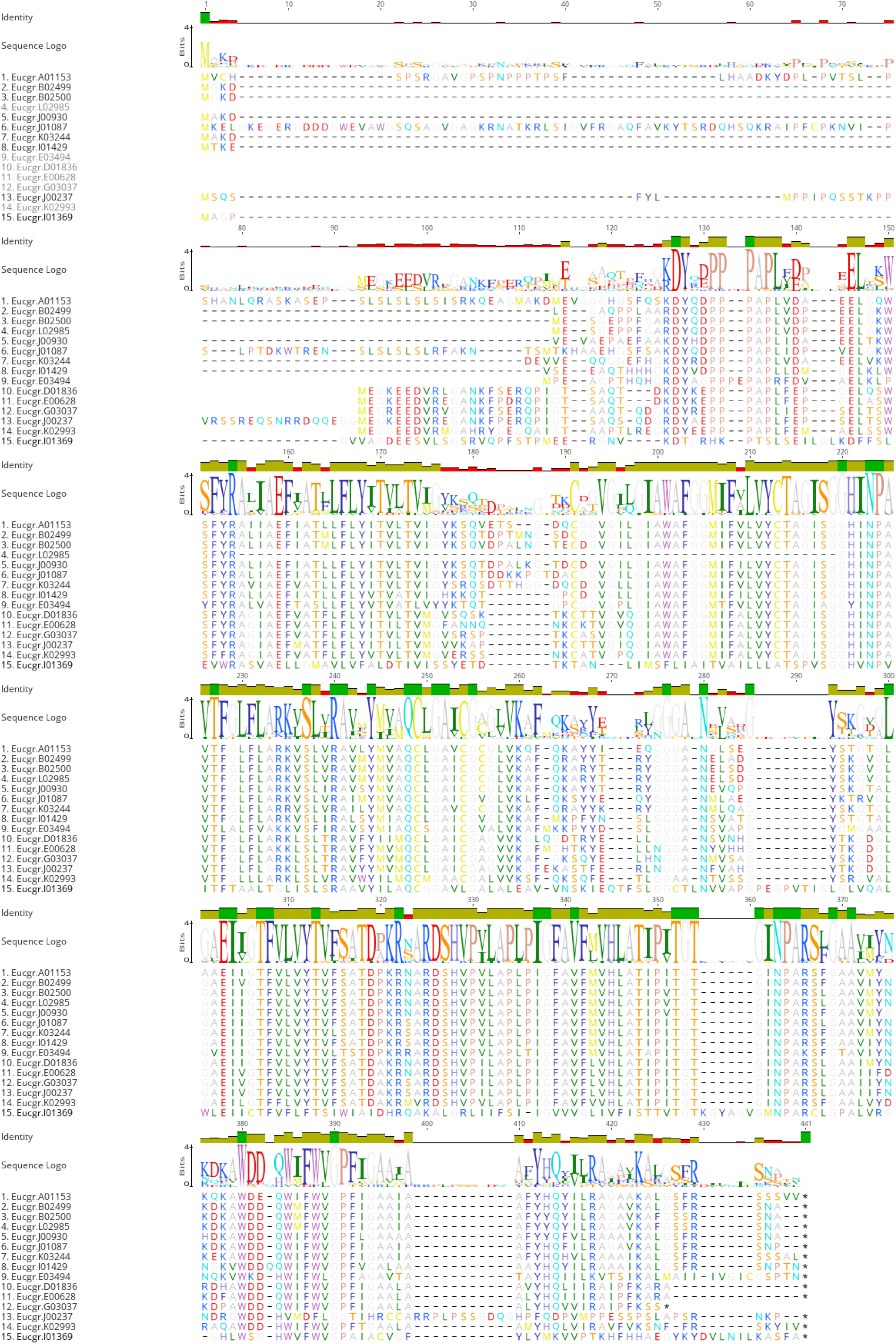
Multiple alignment of EgAQP-PIP proteins.

**Figure S4.**
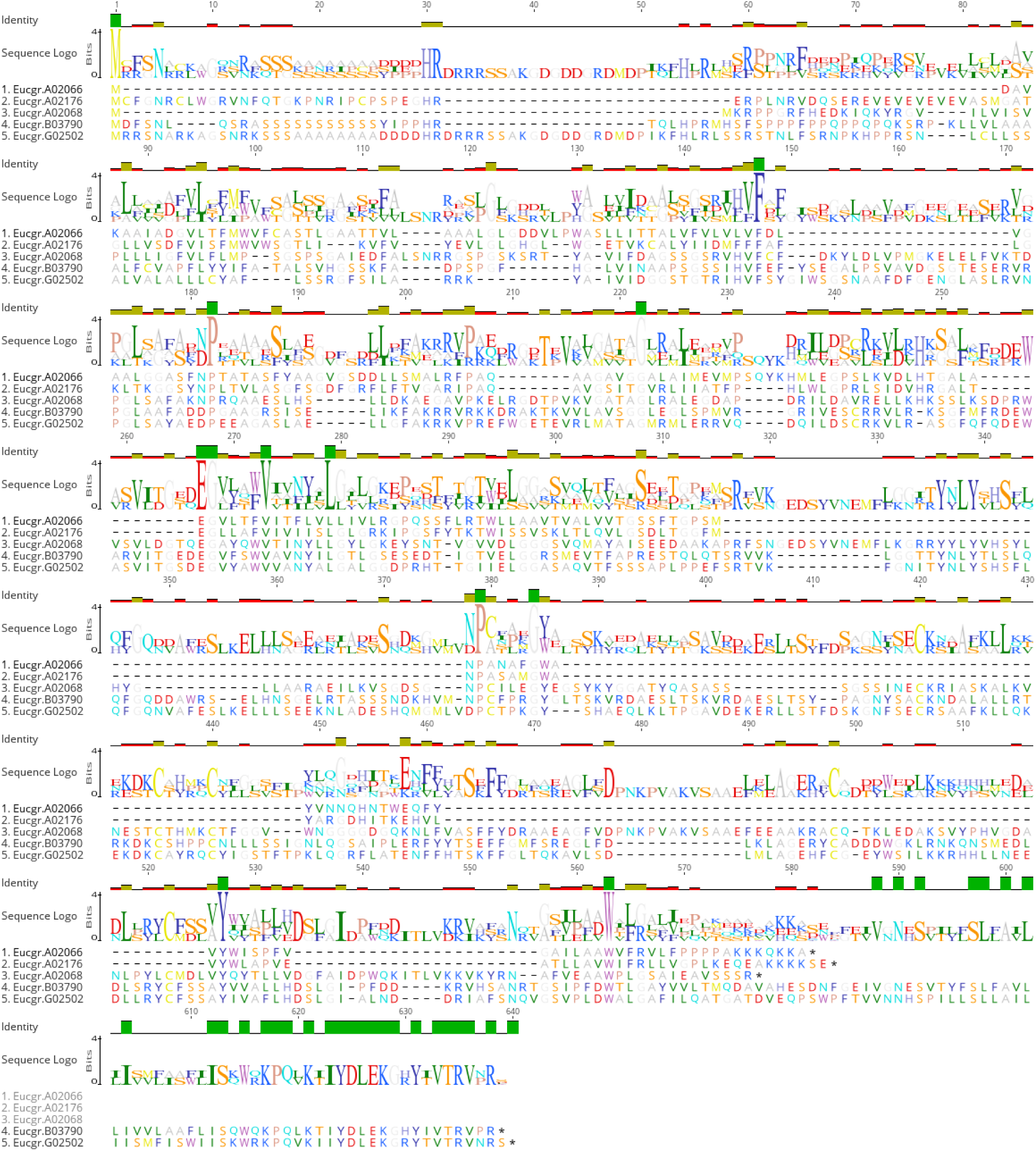
Multiple alignment of EgAQP-SIP proteins.

**Figure S5.**
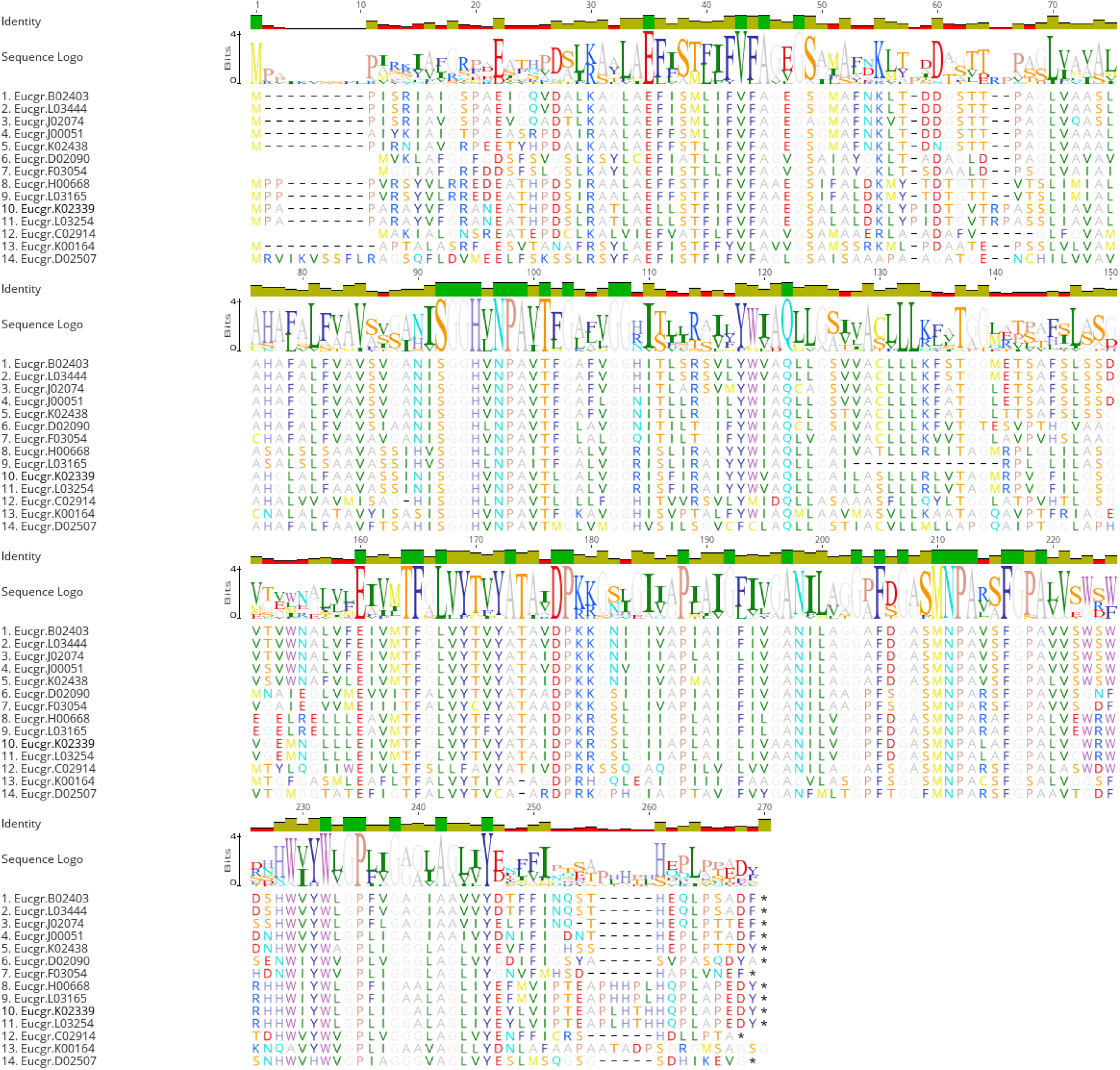
Multiple alignment of EgAQP-NIP proteins.

